# Structural insights into regulation of TRPM7 divalent cation uptake by the small GTPase ARL15

**DOI:** 10.1101/2023.01.19.524765

**Authors:** Luba Mahbub, Guennadi Kozlov, Pengyu Zong, Sandra Tetteh, Thushara Nethramangalath, Caroline Knorn, Jianning Jiang, Ashkan Shahsavan, Emma Lee, Lixia Yue, Loren W. Runnels, Kalle Gehring

## Abstract

Cystathionine-β-synthase (CBS)-pair domain divalent metal cation transport mediators (CNNMs) are an evolutionarily conserved family of magnesium transporters. They promote efflux of Mg^2+^ ions on their own or uptake of divalent cations when coupled to the transient receptor potential ion channel subfamily M member 7 (TRPM7). Recently, ADP-ribosylation factor-like GTPase 15 (ARL15) has been identified as CNNM binding partner and an inhibitor of divalent cation influx by TRPM7. Here, we characterize ARL15 as a GTP-binding protein and demonstrate that it binds the CNNM CBS-pair domain with low micromolar affinity. The crystal structure of the complex between ARL15 GTPase domain and CNNM2 CBS-pair domain reveals the molecular determinants of the interaction and allowed the identification of mutations in ARL15 and CNNM2 mutations that abrogate binding. Loss of CNNM binding prevented ARL15 suppression of TRPM7 channel activity in support of previous reports that the proteins function as a ternary complex. Binding experiments with phosphatase of regenerating liver 2 (PRL2 or PTP4A2) revealed that ARL15 and PRLs compete for binding CNNM, suggesting antagonistic regulation of divalent cation transport by the two proteins.

## Introduction

Magnesium (Mg^2+^) is the most abundant intracellular divalent cation and essential for key cellular processes such as energy production and protein synthesis. In order to maintain and regulate magnesium levels, cells possess a number of Mg^2+^ channels and transporters. Among these are transient receptor potential ion channel subfamily M member 7 (TRPM7) and cystathionine-β-synthase (CBS)-pair domain divalent metal cation transport mediators (CNNMs). TRPM7 is a ubiquitously expressed ion channel with a C-terminal kinase domain also involved in Ca^2+^ homeostasis and Zn^2+^ transport (Duan et al., 2018; Faouzi et al., 2017; Monteilh-Zoller et al., 2003; Runnels et al., 2001; Ryazanova et al., 2010). CNNMs are a widely conserved family of integral membrane proteins and mutated in two genetic diseases linked to Mg^2+^ uptake or transport (Funato and Miki, 2019; Gimenez-Mascarell et al., 2019; Wang et al., 2003).

Recently, TRPM7 and CNNMs were found to function together to mediate divalent cation influx as a trimeric complex with ADP-ribosylation factor-like GTPase 15 (ARL15)(Bai et al., 2021; Kollewe et al., 2021; Zolotarov et al., 2021). Mg^2+^ uptake experiments demonstrated that CNNMs employ the TRPM7 channel and that in the absence of the channel, CNNM2 and CNNM4 can lower intracellular Mg^2+^ levels (Bai *et al*., 2021). Experiments showed ARL15 binds the C-terminal portion of CNNM2 and is required for complex N-glycosylation of CNNM3 (Zolotarov et al., 2021). ARL15 is a member of the RAS superfamily of small GTPases and associated with several metabolic traits including increased risk of diabetes and disorders of lipid metabolism (Richards et al., 2009; Rocha et al., 2017; Wu et al., 2021). While ADP-ribosylation factor (ARF) GTPases are largely involved in membrane-trafficking pathways membrane, the subfamily of ARF-like (ARL) proteins have more diverse functions (Burd et al., 2004; Sztul et al., 2019).

Structurally, CNNMs consist of an N-terminal extracellular domain, a transmembrane domain, and two cytosolic domains: a CBS-pair domain (also termed a Bateman domain) and a cyclic nucleotide-binding homology (CNBH) domain (de Baaij et al., 2012). Two structures of orthologous prokaryotic (CorB) proteins have been determined, confirming that the CNNMs are ion transporters (Chen et al., 2021; Huang et al., 2021). The CBS-pair and CNBH domains from human CNNMs have been extensively studied with multiple structures determined including complexes with phosphatases of regenerating liver (PRLs) (Chen et al., 2018; Chen et al., 2020; Corral-Rodriguez et al., 2014; Gimenez-Mascarell et al., 2017; Gulerez et al., 2016; Zhang et al., 2017). PRL phosphatases regulate Mg^2+^ transport by CNNM proteins and promote of tumor progression and cellular proliferation in cancer (Funato et al., 2014; Hardy et al., 2015).

Here, we used a variety of biophysical techniques to study the interaction of ARL15 with CNNM proteins. Isothermal titration calorimetry (ITC) and NMR experiments demonstrate that ARL15 binds GTP but only with micromolar affinity, orders of magnitude less well than typical GTPases. We show that ARL15 binds CNNM CBS-pair domains with low micromolar affinity and determined the crystal structure of a complex, which shows the ARL15-binding site overlaps with the previously identified PRL-binding site. ITC and NMR experiments confirm that the proteins compete for CNNM binding. The structure also allowed us to design mutants that specifically disrupt ARL15-CNNM binding. Loss of CNNM-binding leads to a loss in the ability of ARL15 to suppress TRPM7 channel activity in HEK-293 cells, confirming the biological relevance of the protein-protein contacts in the crystal structure.

## Results

### ARL15 binds GTP with micromolar affinity

We prepared recombinant ARL15 protein using bacterial expression. Full-length ARL15 showed a propensity to aggregate that hampered its analysis so a truncated version encompassing only the ARL15 GTPase domain (residues 32-197) was prepared. The N-terminal residues removed are the site of ARL15 palmitoylation and form an amphipathic α-helix that promotes membrane binding (Wu *et al*., 2021). The seven residues removed from the C-terminus have no known function. Two-dimensional NMR spectra confirmed that the truncated protein was soluble and well-folded (Figure 1A).

**Figure 1.**
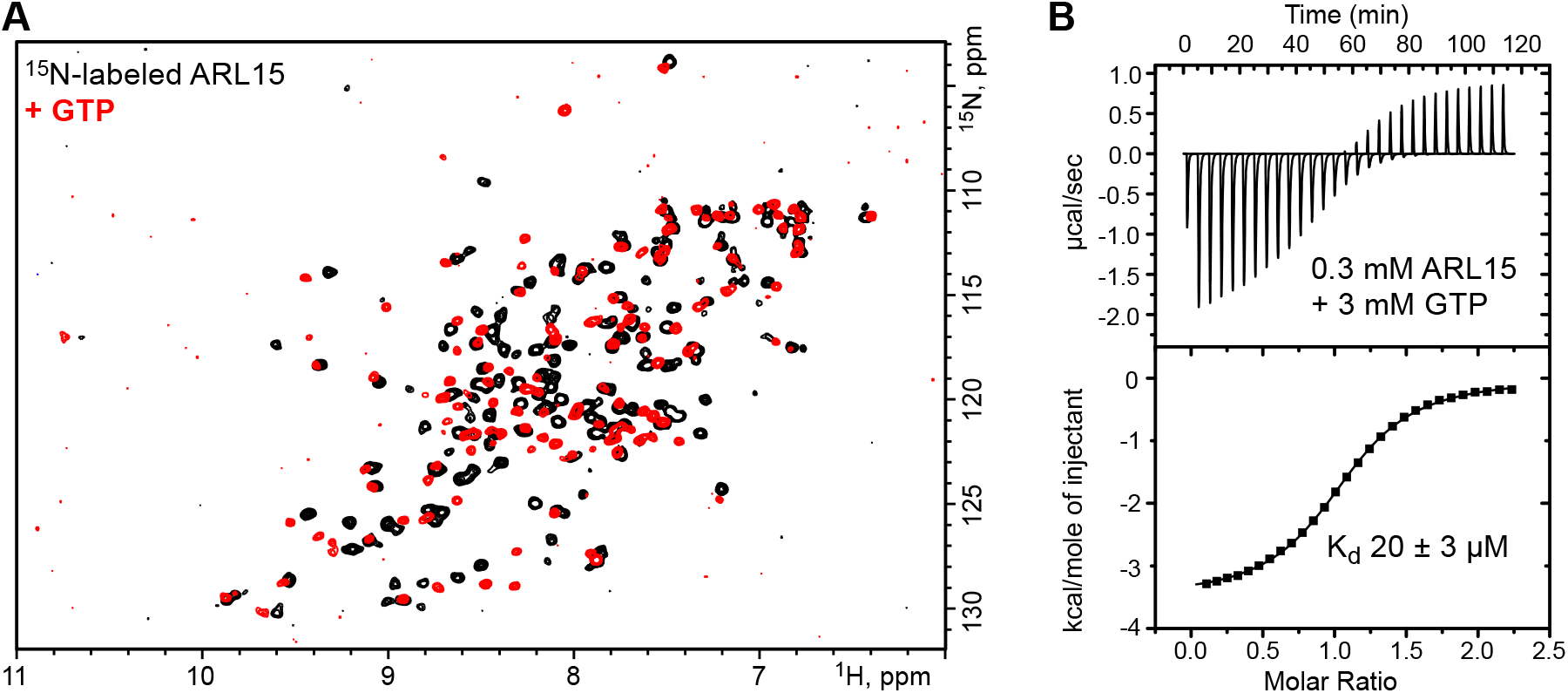
ARL15 has weak affinity for GTP. (**A**) Comparison of ^1^H-^15^N correlation NMR spectra of ARL15 (32-197) unliganded (*black*) and with GTP bound (*red*). (**B**) Isothermal titration calorimetry with ARL15 (32-197) shows it binds GTP with mid-micromolar affinity (N=1.08, K_a_=49,900 M^-1^, ΔH=-3500 cal/mol, ΔS=9.54 cal/mole/deg).

Experiments to preload ARL15 with GTP or GDP were unsuccessful, and we observed that the protein purified from *E. coli* was nucleotide-free. NMR spectra showed addition of GTP induced many changes with a general improvement in the uniformity and intensity of the signals (Figure 1A). To measure the affinity of ARL15 for GTP, we used isothermal titration calorimetry (ITC). Experiments at low protein concentrations showed only a small signal but at 300 μM ARL15, a clear and unambiguous thermogram was observed (Figure 1B). Fitting the data with a single binding site model, we measured an affinity of 20 μM, almost six orders of magnitude weaker than the picomolar affinity of other small GTPases, such as Ras (Ford et al., 2009). Attempts to detect GTPase activity were unsuccessful either due to the absence of a GTPase activating protein (GAP) or an intrinsic lack of catalytic activity.

### ARL15 binds CNNMs with low micromolar affinity

To quantify the interaction between ARL15 and CNNM2, we prepared constructs containing the CBS-pair domain (residues 429-584) and a larger cytosolic fragment including the cyclic-nucleotide binding homology (CNBH) domain (Figure 2A). When titrated with ARL15 (32-197), both constructs generated a strong exothermic signal, corresponding to binding in the range of 1 to 2 μM (Figure 2B, Suppl. Figure S1). We did not observe significant increase in affinity when the CNBH domain was present in disagreement with a previous co-immunoprecipitation study (Zolotarov *et al*., 2021).

**Figure 2.**
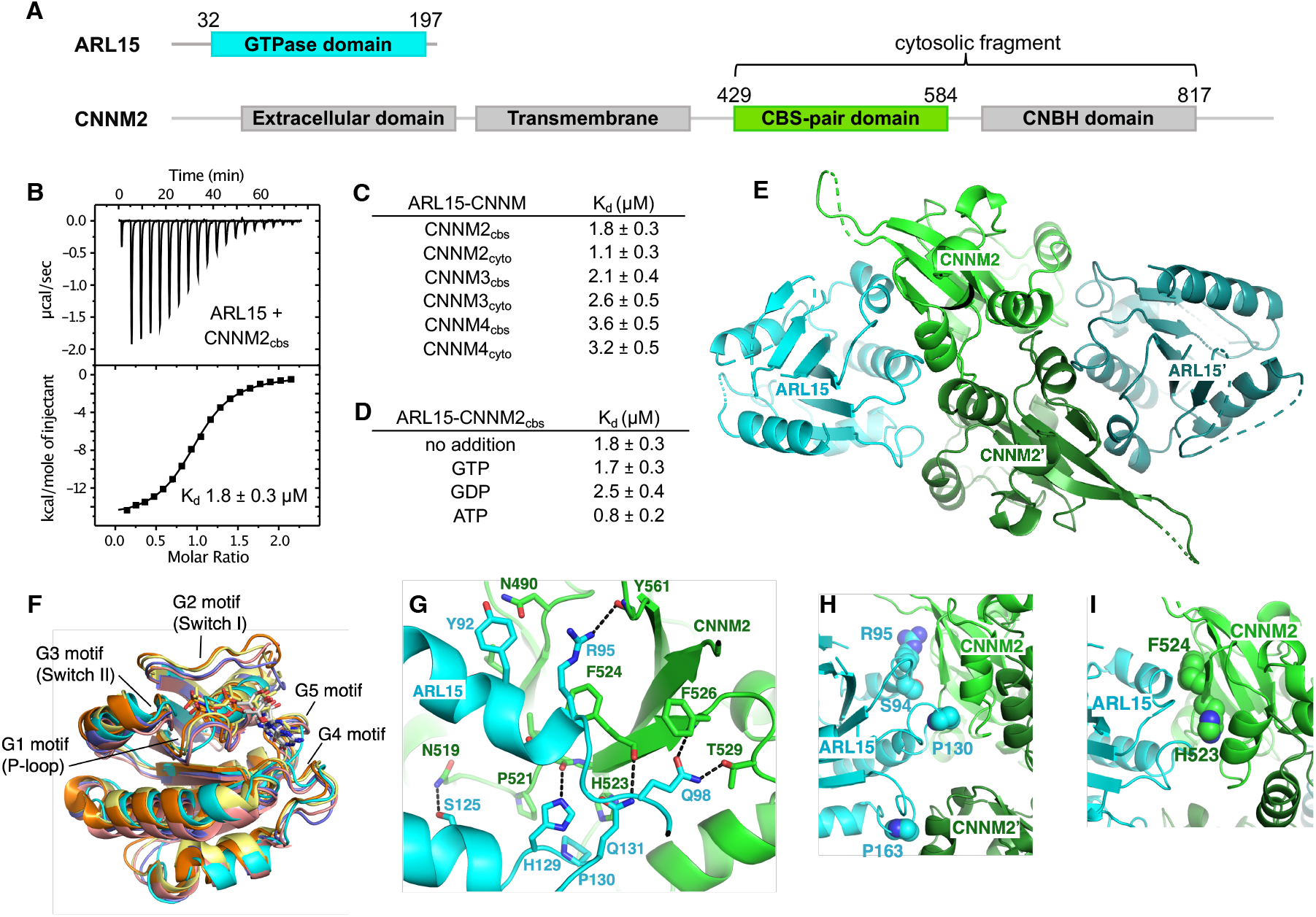
ARL15 binds CNNM CBS-pair domains. (**A**) Domain organization of ARL15 and CNNM proteins. All four CNNM protein have the same domain organization but differ in size. (**B**) Isothermal titration calorimetry experiment between the ARL15 GTPase domain and CNNM2 CBS-pair domain. (**C**) Binding affinities (K_d_) between ARL15 GTPase domain and CBS-pair and cytosolic fragments of CNNM2, CNNM3 and CNNM4 measured by ITC. (**D**) Effect of nucleotides on the affinity of the ARL15-CNNM2 interaction measured by ITC. (**E**) Crystal structure of ARL15 GTPase domain bound to the CBS-pair domain of CNNM2. Two ARL15 molecules bind symmetrically to the CBS-pair domain dimer. (**F**) Structural comparisons of ARL15 to other GTPases: ARL2 (*pink*, PDB 4GOK), ARL3 (*orange*, PDB 3BH6), ARF1 (*yellow*, PDB 1O3Y), ARF6 (*violet*, PDB 4KAX). (**G**) Detail of contacts between ARL15 and one of the monomers in the CNNM2 dimer. (**H**) Contacts between ARL15 and the two monomers in the CNNM2 dimer. ARL15 residues selected for mutagenesis are shown. (**I**) CNNM2 residues selected for mutagenesis.

Experiments with the corresponding fragments of CNNM3 and CNNM4 also showed low micromolar affinities, demonstrating that ARL15-binding is a conserved property of CNNM CBS-pair domains (Figure 2C). We next tested whether nucleotide-binding affects the interaction. Unexpectedly, CNNM binding was not affected by the absence or presence of bound GTP or GDP, suggesting that the ARL15 nucleotide-binding switch regions are not involved (Figure 2D). We also tested if Mg^2+^-ATP affects ARL15 binding. Mg^2+^-ATP binds to CNNM CBS-pair domains as promotes dimerization in a flat ring conformation (Corral-Rodriguez *et al*., 2014). We observed no change in ARL15 affinity when Mg^2+^-ATP was present in agreement with co-immunoprecipitation experiments (Zolotarov *et al*., 2021). This suggests that CBS-pair dimerization does not regulate ARL15-binding.

### Structure of the ARL15-CNNM CBS-pair domain complex

We turned to X-ray crystallography to visualize the ARL15-CNNM interaction. Commercial crystallization screens were used for mixtures of ARL15 (32-197) with cytosolic or CBS constructs of CNNM2, CNNM3 and CNNM4. Small crystals were obtained with a sample containing ARL15 and CNNM2 CBS-pair domain (429-584). The crystals were improved by moving the His-tag on ARL15 to the C-terminus. The best crystals diffracted to 3.2 Å using synchrotron radiation. The structure was solved by molecular replacement using CNNM2 CBS structure (PDB 4IY0) and the AlphaFold model of ARL15. The large asymmetric unit contains four ARL15 and four CNNM2 CBS-pair domains. The relative position of the ARL15 molecules to CBS-pair domains was well defined with 0.6 Å RMSD between the four copies. The protein interfaces were among the best-defined areas in the electron density map. Outside of these sites, there were deviations in the conformations of some loops and the orientation of the CBS-pair domain N-terminal helix, which are likely due to crystal packing (Figure 2E).

### Structural comparisons of ARL15 to other GTPases

Comparison of the ARL15 structure with other GTPases confirmed it belongs in the ARF/ARL family (Figure 2F). An RMSD of ~1.5 Å was observed between ARL15 and the GTPases ARL2, ARL3, and ARF6, reflecting the high degree of sequence identity (around 35%). Unlike structures of most GTPases, ARL15 showed a large degree of disorder, not only in loops, but also in the GTP-binding site. The G2 and G5 motifs were largely disordered. The G1 motif (P-loop) was traceable but adopted a conformation incompatible with nucleotide binding. Disorder in the absence of a bound nucleotide could also be seen in the NMR spectra where, upon addition of GTP, many signals became sharper and more intense (Figure 1A).

### Identification of the ARL15-CNNM interface

In the crystal, the ARL15 makes contacts with two CBS-pair domains (Figure 2E). The CBS-pair domains are present as a dimer but compared to previous structures, the dimer is twisted, possibly due to the absence of Mg^2+^-ATP. This twist opens the dimer slightly generating two contact surfaces between ARL15 and the CNNM2 CBS-pair domains. The larger surface contains numerous polar contacts and involves two structural elements of ARL15: a C-terminal part of helix α2 with following loop and the loop following helix α3 (Figure 2G). Helix α2 is a continuation of the Switch II region and is a common binding site for many GTPase effectors. The side chain of ARL15 Gln98 makes intermolecular hydrogen bonds with side chain of Thr529 and backbone amide of Phe526. Along the same β-strand of CBS domain, side chains of Gln131 and His129 form hydrogen bonds with backbone carbonyl groups of Phe524 and Leu522, respectively. In one of the copies of the ARL15-CBS complex, the side chain of Arg95 bonds with backbone carbonyl of Tyr561. The side chain of Arg95 appears to be partially disordered in other copies which could be a result of competing interactions with other nearby electron donors such as a mainchain carbonyl or the side chain of Asn490. Significantly, all the ARL15-CNNM2 interfaces show hydrophobic stacking between aliphatic part of Arg95 and the side chain of Phe524. Another key hydrophobic binding determinant is provided by ARL15 Pro130 that inserts into a small pocket formed by side chains of CNNM2 Pro521 and His523.

The second binding surface between ARL15 and the CBS-pair domain is smaller and involves ARL15 residues in the loop following helix α4 that sit in a shallow pocket formed by two helices from the CBS-pair domain (Figure 2E). These helices are usually part of the CBS-pair dimerization interface and not exposed in the flat dimer typically observed in the presence of Mg^2+^-ATP.

### Identification of the interaction surface by mutagenesis

We turned to mutagenesis to determine the importance of the two binding surfaces. We mutated Ser94, Arg95 and Pro130 of ARL15 from the larger interaction surface and Pro163 from smaller interface (Figure 2H). We confirmed that the mutations did not cause unfolding of ARL15 using NMR spectroscopy. 1D spectra of the mutants closely matched the spectrum of the wild-type protein (Suppl. Figure S2). We tested the mutant proteins for binding to CBS-pair domain of CNNM2 (residues 429-584) using pulldown assays and ITC. Pulldowns with GST-fused CNNM2 CBS-pair domain showed binding for wild-type ARL15 and the P163K mutant but not for S94K, R95A and P130W (Figure 3A). Identical results were obtained in pulldowns using GST-fused CBS-pair domains of CNNM3 and CNNM4 (Figure 3B & 3C). The pulldown results were confirmed by ITC experiments with the CNNM2 CBS-pair domain. The S94K and S94W mutations decreased the affinity of ARL15 towards CBS-pair domain by 20-fold (K_d_ of 40 μM), while the R95A and P130W mutations abrogated the binding (Figure 3D & 3E). On the other hand, the P163K and P163W mutants showed wild-type affinity. These results demonstrate that the first interaction surface is critical for the ARL15-CNNM binding in solution. The second surface is dispensable and likely the result of crystal packing.

**Figure 3.**
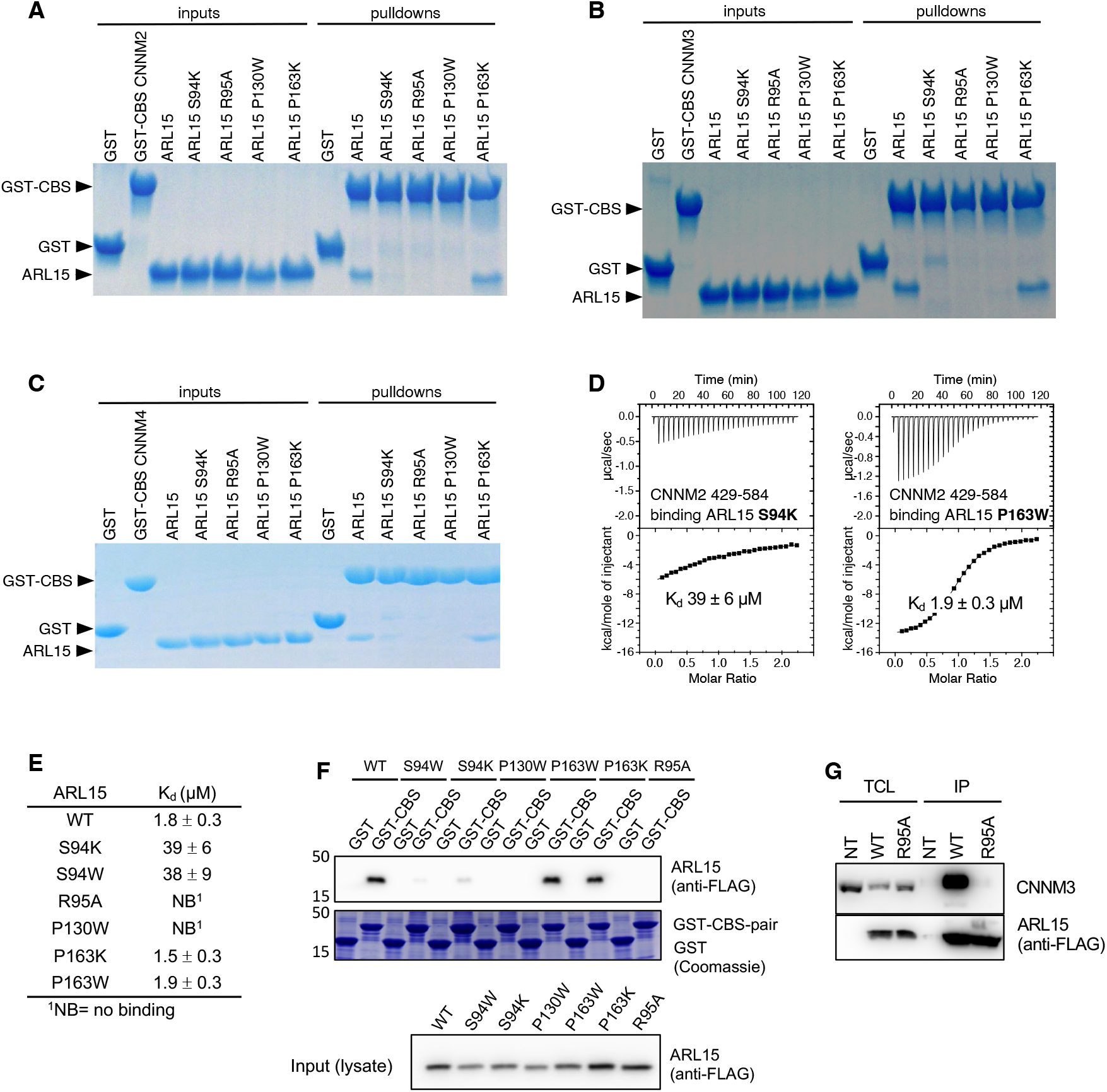
Mutagenesis of ARL15 confirms the CNNM-binding site. Pulldowns of recombinant WT and mutant ARL15 GTPase domains (residues 32-197) by (**A**) GST-CNNM2 CBS-pair domain (residues 429-584), (**B**) GST-CNNM3 CBS-pair domain (residues 299-452), (**C**) GST-CNNM4 CBS-pair domain (residues 356-511). (**D**) Isothermal titration of mutants of ARL15 GTPase domain and CNNM2 CBS-pair domain. (**E**) Binding affinities (K_d_) of ARL15 GTPase mutants measured by ITC. (**F**) Pulldown of wild-type ARL15 (WT) and the indicated mutants expressed in HEK-293T cells by GST-fused CNNM2 CBS-pair domain. (**G**) Co-immunoprecipitation of FLAG-tagged ARL15 with native CNNM3 shows the R95A mutation fully blocks binding. NT, non-transfected; WT, wild-type.

We confirmed the results in HEK-293T cells using a GST-pulldown and co-immunoprecipitation experiments. GST-tagged CBS-pair domain of murine CNNM2 strongly bound to FLAG-tagged full-length human ARL15 (Figure 3F). The P163W and P163K mutants showed wild-type binding. Weak binding was observed with S94W and S94K and no binding observed with the P130W and R95A mutants. The same interaction surface is essential for binding full-length CNNM. HEK293T cells were transfected with wild-type and R95A mutant FLAG-tagged ARL15 to analyze the binding interaction with endogenous CNNM. Native CNNM3 was co-immunoprecipitated with FLAG-tagged ARL15 wild-type (WT) but not with FLAG-tagged ARL15 mutant (R95A) or non-transfected (NT) cells (Figure 3G). It indicates that the mutation of Arg95 is sufficient to disrupt the binding of the entire protein.

### The ARL15-binding site is conserved across the CNNM family

We used the crystal structure to identify point mutations in CNNMs that would knock out ARL15 binding (Figure 2I). Residues H523 and F524 in CNNM2 correspond to H391 and F392 in CNNM3 and H450 and F451 in CNNM4. Mutations in the CBS-pair and cytosolic fragments of the three CNNM proteins were prepared, and proper folding of the mutants confirmed by NMR spectroscopy (Suppl. Figure S3). The binding to ARL15 was measured in pulldown and ITC experiments (Figure 4). In all cases, the mutations caused a complete loss of ARL15 binding in the pulldown assays with either the CBS-pair domains or cytosolic fragments. ITC titrations appeared to be more sensitive to weak binding, and a small amount of residual binding affinity was observed with the CNNM2 F524, CNNM3 F392 and CNNM4 F451 mutations (Figure 5E, Suppl. Figure S1). In contrast, the loss of the histidine (H523, H391, or H450) led to a complete loss of signal in the ITC experiments. The concurrence of the mutagenesis results across the three CNNM isoforms confirms that ARL15 binds identically to all three proteins.

**Figure 4.**
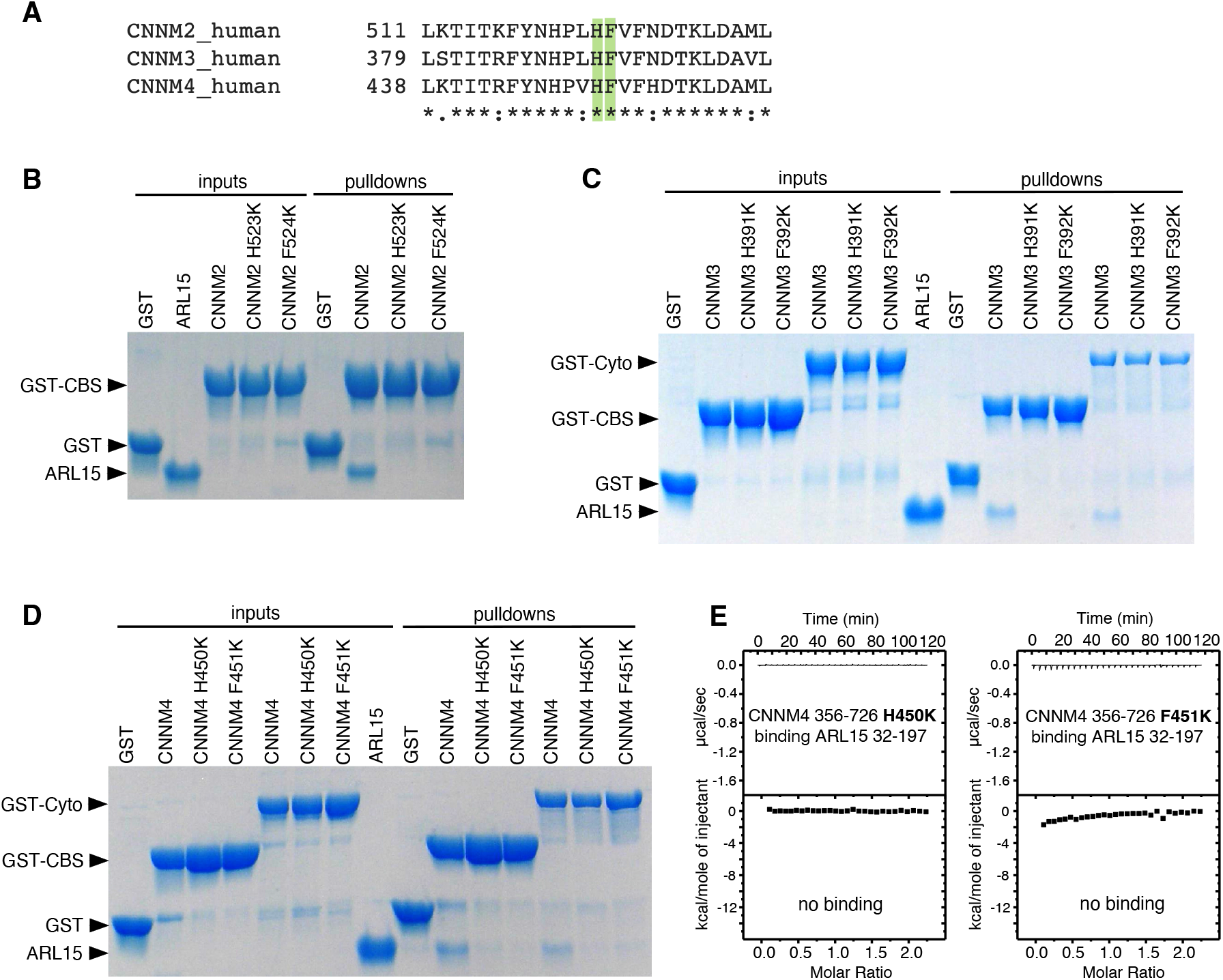
Mutagenesis of CNNM confirms conservation of ARL15-binding site. (**A**) Conservation of ARL15-binding residues in all three CNNMs. (**B**-**D**) Pulldown of recombinant GTPase domain (residues 32-197) by (**B**) WT and mutant GST-CNNM2 CBS-pair domain (residues 429-584), (**C**) WT and mutant GST-CNNM3 CBS-pair domain (residues 299-452) and cytosolic fragment (residues 299-658). (**D**) WT and mutant GST-CNNM4 CBS-pair domain (residues 356-511) and cytosolic fragment (residues 356-726). (**E**) Isothermal titration of ARL15 GTPase domain and two mutants of the CNNM4 cytosolic fragment.

**Figure 5.**
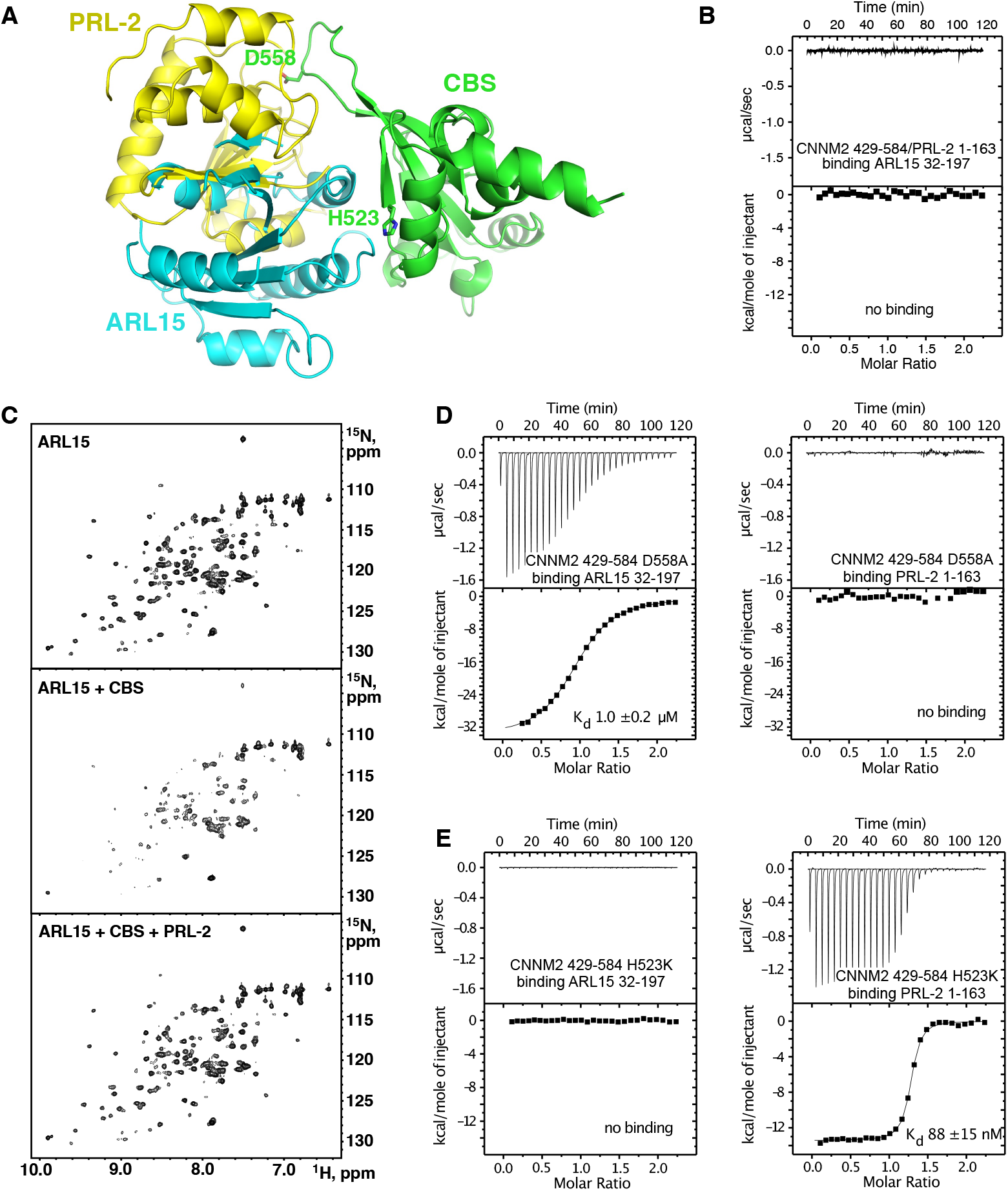
ARL15 and PRL2 have overlapping but distinct binding sites. (**A**) Overlay of the structures of the CNNM2 complexes with ARL15 (*cyan*) and PRL2 (*yellow*) shows simultaneous binding is not possible. (**B**) ITC experiment shows no binding of ARL15 to the preformed complex of CNNM2 and PRL2. (**C**) ^1^H-^15^N correlation NMR spectra of _15_N-labeled ARL15 alone (top), in the presence of the CNNM2 CBS-pair domain (middle), and after addition of PRL2 (bottom). The ARL15 NMR signals are attenuated upon binding CNNM2 but reappear when ARL15 is displaced by PRL2. (**D**) ITC experiments demonstrating that the CNNM2 D558A mutation specifically disrupts PRL2 binding. (**E**) ITC experiments demonstrating that the CNNM2 H523K mutation specifically disrupts ARL15 binding.

### ARL15 and PRL phosphatases compete for binding

Both ARL15 and PRL phosphatases bind CNNM; therefore, it is relevant to ask if they compete for binding. Structural superposition the complexes with the CNNM2 CBS-pair domain shows considerable overlap in positions of ARL15 and PRL2 (Figure 5A). This strongly suggests they cannot bind at the same time. We used ITC to test for competitive binding. PRL phosphatases bind to CNNM with approximately 100-fold better affinity and would be expected to outcompete ARL15 for binding. Indeed, titration of ARL15 into a mixture of CNNM2 CBS and PRL2 resulted in essentially no heat released indicating no binding occurred (Figure 5B). In the cell, competition between the two proteins will be affected by relative abundance of ARL15 and PRLs and by the regulation of PRL binding. When the catalytic cysteine of PRL is modified via oxidation or phosphorylation, the affinity of PRL for CNNMs is decreased or abolished, potentially upregulating ARL15 binding (Gulerez *et al*., 2016). Note that the competition does not prevent ARL15 and PRLs binding simultaneously to separate monomers of a CNNM dimer.

As independent verification, we designed a competition experiment using NMR (Figure 5C). Addition of the CNNM2 CBS-pair domain (429-584) to ^15^N-labeled ARL15 (32-197) results in disappearance of a number of signals in the ^1^H-^15^N correlation spectrum due to the formation of a higher molecular weight complex. In contrast, addition of PRL2 with the CBS-pair domain did not change in the spectrum due to PRL2 outcompeting ARL15 for binding.

### Partner specific CNNM mutants

Analysis of the structures suggested it should be possible to design CNNM mutants that specifically block binding of ARL15 or PRLs. We used ITC to test the ability of D558A and H523K mutants of the CNNM2 CBS-pair domain for binding ARL15 and PRL2 (Figure 5D & E). Aspartic acid 558 was previously shown to be required for CNNM3 binding PRL1 {Gimenez-Mascarell, 2017 #216}. The D558A mutant bound to ARL15 with the same affinity as wild-type CBS-pair domain, while it was unable to interact with PRL2. Conversely, the H523K mutant bound PRL2 with nanomolar affinity but completely blocked ARL15 binding. These mutants should be useful in future biological studies since they differentiate ARL15 and PRL binding.

### CNNM-binding-defective ARL15 is unable to inhibit TRPM7 channel activity

Overexpression of ARL15 has been reported to inhibit Zn^2+^ influx in cells by TRPM7 (Bai *et al*., 2021; Kollewe *et al*., 2021). To test the importance of the ARL15-CNNM interaction, we examined the effects of transfection of wild-type ARL15 and the R95A mutant in two assays of TRPM7 channel function. Whole-cell electrophysiological recordings of cells with and without co-expression of ARL15 confirmed that ARL15 suppresses ion influx (Figure 6A & B). In contrast, co-expression of the ARL15 R95A mutant showed no suppression with currents indistinguishable from the non-transfected cells. Similar results were observed with a Zn^2+^-influx assay, taking advantage of TRPM7 permeability to Zn^2+^ (Monteilh-Zoller *et al*., 2003), comparing cells with (293-TRPM7) and without (HEK-293T) expression of TRPM7 (Figure 6C & D). ARL15 co-expression markedly decreased intracellular Zn^2+^ levels in TRPM7-expressing cells while expression of the ARL15 R95A mutant had no effect.

**Figure 6.**
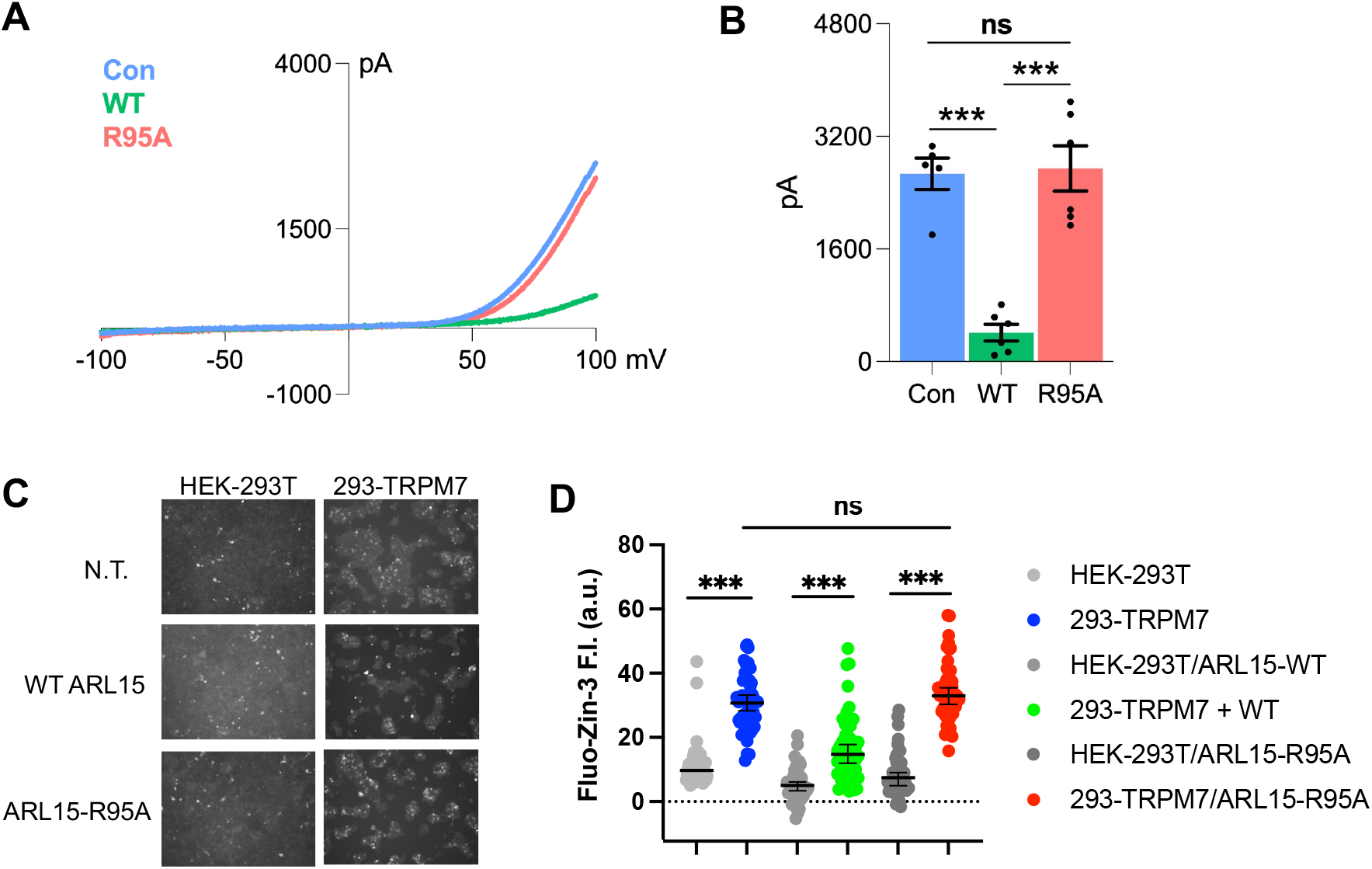
Disruption of the CNNM-binding site blocks ARL15 regulation of TRPM7 Zn^**2+**^ **influx**. (**A**) Representative TRPM7 whole cell currents from 293-TRPM7 cells from non-transfected controls (Con), wild-type ARL15 (WT), and ARL15 R95A mutant (R95A) transfectants. (**B**) Average current density of the different groups from (A). *** indicates a *p*-value of less than 0.0001. (**C**) Zinc influx assay using the FluoZin-3 Zn^2+^ indicator was used to monitor TRPM7 function in non-transfected (N.T.) cells or transfected with wild-type or mutant ARL15. Images shown are taken at a time point 5 to 10 minutes after application of 30 μM ZnCl_2_ to stimulate Zn^2+^ influx. (**D**) Cell counts of fluorescence intensity (F.I.) from (C). A total of 50 cells per condition were randomly selected for quantification. *** indicates a *p*-value of less than 0.0001.

## Discussion

### ARL15 is an atypical GTPase

ARL15 has been identified in genome-wide association studies of coronary heart disease, kidney disease, rheumatoid arthritis, and diabetes (Gorski et al., 2017; Mahajan et al., 2014; Negi et al., 2013). Electrophysiological studies have shown that ARL15 inhibits ion transport through the channel kinase TRPM7 (Bai *et al*., 2021; Kollewe *et al*., 2021). The weak affinity for GTP is a property shared with several other ARL-family members. ARL2, ARL3 and ARL13B all have been reported to bind guanine nucleotides with micromolar affinity (Hanzal-Bayer et al., 2005; Ivanova et al., 2017; Linari et al., 1999). In this affinity range, it is unclear if a GEF is required to facilitate GTP binding.

We see rapid binding of GTP to nucleotide-free in ITC experiments (Figure 1B). Similarly, a GAP would seem to be unnecessary since the bound nucleotides exchange rapidly. We did not observe intrinsic GTPase activity, which could be related to the presence of an alanine residue at position 86 (Suppl. Figure S4). In other ARLs, this position is occupied by glutamine, which plays a role in GTP hydrolysis and is often mutated to leucine to block GTPase activity (Sprang, 1997; Sztul *et al*., 2019). In the presence of rapid nucleotide exchange, GTP hydrolysis would generate a wasteful, futile cycle. A second notable sequence difference is the connecting region between Switch I and Switch II that is characterize of ARF/ARL GTPase family members. This region contains two β-strands and a patch of three conserved aromatic residues that participate in the interactions with effectors (Menetrey et al., 2007). In ARL15, the three residues are Phe66, Lys81 and Tyr96. Lys81 replaces a tryptophan found in all ARFs and many ARLs, which suggests that ARL15 might interact with effectors differently from other ARF/ARL GTPases.

### Complex with CNNM

CBS-pair domains, also termed Bateman domains, are found in many proteins including a chloride channel ClC and bacterial Mg^2+^ transporter MgtE (Bateman, 1997; Baykov et al., 2011). They consist of repeated CBS motifs - CBS1 and CBS2 - that together form a Mg^2+^-ATP binding site (Ereno-Orbea et al., 2013). The CNNM CBS-pair domains most often dimerize in a head-to-head manner forming a ring around two ATP molecules (Corral-Rodriguez *et al*., 2014). Considerable plasticity exists at the dimerization interface with many different conformations observed in crystal structures (Chen *et al*., 2020; Chen *et al*., 2021; Gimenez-Mascarell *et al*., 2017; Gulerez *et al*., 2016; Zhang *et al*., 2017).

Our structure of the complex of ARL15 bound to the CNNM2 CBS-pair domain differs from a previously proposed *in silico* model (Zolotarov *et al*., 2021). The model suggests that the first CBS motif and CNBH domain of CNNM2 bind ARL15. This is inconsistent with our observations that residues, His523 and Phe524, in the second CBS motif are essential for binding, and deletion of the CNBH domain has no effect. The model also predicts the wrong face of ARL15 with only peripheral involvement of Arg95 and Pro130.

In addition to ARL15, CNNM CBS-pair domains bind PRLs with low nanomolar affinity (Gulerez *et al*., 2016). Binding is mediated by an aspartic acid that inserts into the phosphatase catalytic site, mimicking a phosphoprotein substrate (Gehring et al., 2022; Gimenez-Mascarell *et al*., 2017; Gulerez *et al*., 2016). While the binding sites overlap, a CNNM dimer could simultaneously bind ARL15 and PRLs using separate CBS-pair domains. Based on its ~100-fold higher affinity *in vitro*, PRLs should outcompete ARL15 for CNNM binding but this would depend on the protein concentrations. Little is known about the relative concentrations of ARL15 and PRLs or their localization in the cell. Endogenous ARL15 is reversibly palmitoylated and has been reported to be dynamically localized (Wu *et al*., 2021). PRLs are farnesylated and membrane bound. They undergo reversible modifications of their catalytic cysteine, which block CNNM binding, and could act to upregulate ARL15 binding.

### Cellular function

ARL15 has been reported to form a complex with CNNMs and TRPM7 (Kollewe *et al*., 2021; Zolotarov *et al*., 2021). While it has not been reported to affect CNNM activity, two groups reported that it inhibits divalent cation uptake by TRPM7 (Bai *et al*., 2021; Kollewe *et al*., 2021) and we show that CNNM-binding is required for the inhibition (Figure 6). Independently PRL-binding has been shown to inhibit Mg^2+^ efflux by CNNM4 (Funato *et al*., 2014; Gulerez *et al*., 2016; Kozlov et al., 2020). This leads us to propose a model of regulation of divalent cation transport through competitive binding of PRLs and ARL15 to a complex of CNNM and TRPM7 (Figure 7). While speculative, the model is useful as recapitulates several observations and develops hypotheses for future testing. Among the observations are: 1) PRL overexpression stimulates TRPM7 channel function in the presence of CNNMs {Bai, 2021 #1713}, 2) PRLs inhibit CNNM ion efflux, and 3) CNNM-binding by ARL15 is required for its inhibition of TRPM7 (Figure 6). The model suggests that ARL15-binding may stimulate CNNM activity. This is untested but reasonable given that ARL15 can displace PRLs from CNNMs. As CNNMs are electroneutral ion antiporters (Chen *et al*., 2021; Yamazaki et al., 2013), detection of their activity in electrophysiological experiments is more difficult than detecting TRPM7 activity. Post-translational modifications of ARL15 and PRLs likely play important roles in adjusting the balance of their opposing roles in maintaining magnesium and other divalent cations levels in the cell. In future studies, the identification of CNNM mutations that specifically block PRL and ARL15 binding should clarify their contrasting roles in divalent cation transport.

**Figure 7.**
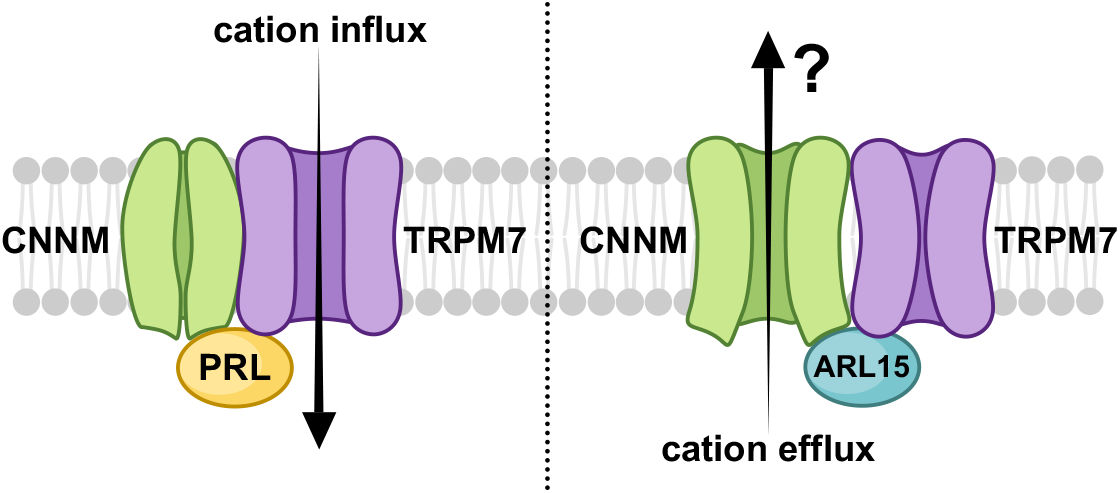
Model of regulation of CNNM-TRPM7 complex by ARL15 and PRLs. PRL binding to CNNM inhibits CNNM activity and activates cation influx via TRPM7. Conversely, ARL15 binding to CNNM inhibits TRPM7 activity. It is unknown if ARL15 binding stimulates CNNM efflux and if efflux occurs in the complex with TRPM7.

## Materials and Methods

### Expression and purification of recombinant proteins

A plasmid for bacterial expression of human ARL15 (1-204) was obtained by cloning codon-optimized synthetic DNA into the NdeI and BamHI cloning sites of pET15b (Bio Basic). A truncated construct (residues 32-197) and point mutants were obtained by mutagenesis using the QuikChange Lightning Site-Directed Mutagenesis Kit (Thermo Fisher Scientific). For crystallization, the N-terminal His-tag was deleted and the HHHHHH sequence was inserted after residue 197 followed by a stop codon. For HEK293T expression, human ARL15 (NM_019087.3) in the pcDNA3.1+/C-(K)-DYK vector (GenScript) was used and further employed to introduce the mutations into the construct. Plasmids expressing human CNNM2 (residues 429-584 and 429-817), CNNM3 (residues 299-452 and 299-658), CNNM4 (residues 356-511 and 356-726) were described previously (Chen *et al*., 2020). The point mutants of CNNMs were obtained by mutagenesis using the QuikChange Lightning Site-Directed Mutagenesis Kit (Thermo Fisher Scientific). The amino acid sequence of the CBS-pair domain of murine CNNM2 (495-582) was codon optimized for bacterial expression and subcloned into the EcoRI/XhoI sites of pGEX-6P3 vector (GenScript). The plasmid expressing His-tagged human PRL2 phosphatase (residues 1–163) was described previously (Gulerez *et al*., 2016). DNA sequencing was used to verify all sequence modifications.

Proteins were expressed in *E. coli* BL21(DE3) at 37 ºC in Luria Broth (LB). ARL15 expression was induced with 0.5 mM IPTG for 4 hours at 30 ºC. CNNM cytosolic fragments were induced with 1 mM IPTG overnight at 18 ºC, while CBS-pair domains were induced with 1 mM IPTG for 4 hours at 30 ºC. For NMR experiments, the recombinant protein was isotopically labeled by growth of *E. coli* BL21 in M9 minimal medium with ^15^N-ammonium sulfate as the sole source of nitrogen.

For His-tagged protein purification, cells were harvested and broken in lysis buffer (50 mM HEPES pH 7.6, 0.5 M NaCl, 5% glycerol) containing 1 mM PMSF, 0.1 mg/ml lysozyme, 0.04% β-mercaptoethanol. His-tagged proteins were purified by affinity chromatography on Ni-NTA Agarose resin (Qiagen) and eluted with buffer containing 0.5 M imidazole. For GST-tagged protein purification, cells were harvested and broken in 1X PBS (phosphate buffered saline) containing 1 mM PMSF, 0.1 mg/ml lysozyme, 0.04% β-mercaptoethanol. GST-tagged proteins were purified on Glutathione Sepharose resin (GE Healthcare) and eluted with buffer containing 20 mM reduced glutathione. The GST-tag was removed by overnight incubation with PreScission Protease, leaving an N-terminal Gly-Pro-Leu-Gly-Ser extension. For final purification, the proteins were applied to a HiLoad 16/600 Superdex 75 or 200 size exclusion column (Cytiva) in HPLC buffer (50 mM HEPES pH 7.5, 200 mM NaCl, 1 mM MgCl_2_, 1 mM tris(2-carboxyethyl)phosphine hydrochloride (TCEP-HCl)). PRL2 (1-163) was purified using HPLC buffer with higher concentration of the reducing agent (50 mM HEPES pH 7.5, 200 mM NaCl, 1 mM MgCl_2_, 5 mM TCEP-HCl). The final purified proteins were concentrated to around 10-25 mg/ml (estimated by UV absorbance), and the purity was verified by SDS-PAGE.

### NMR spectroscopy

NMR samples were 100 μM in 50 mM HEPES pH 7.5, 200 mM NaCl, 1 mM MgCl_2_, 1 mM TCEP-HCl. ^1^H-^15^N correlation experiment was done using 150 μM PRL2 (1-163) in 50 mM HEPES pH 7.5, 200 mM NaCl, 1 mM MgCl_2_, 5 mM TCEP-HCl. All NMR experiments were performed at 25 ºC on a Bruker 600 MHz spectrometer. NMR spectra were processed using NMRPipe (Delaglio et al., 1995) and analyzed with SPARKY (Goddard and Kneller, 2008).

### Isothermal titration calorimetry

ITC experiments were performed on MicroCal VP-ITC titration calorimeter (Malvern Instruments Ltd). The syringe was typically loaded with 150 or 300 μM concentration of the ligand, while the sample cell contained 15 or 30 μM protein. All experiments were carried out at 20 ºC with 19 injections of 15 μl or 29 injections of 10 μl. Results were analyzed using ORIGIN software (MicroCal) and fitted to a binding model with a single set of identical sites.

### Crystallization

Initial crystallization conditions were identified utilizing hanging drop vapor diffusion with the Classics II and ProComplex screens (QIAGEN). The best crystals were obtained by equilibrating a 0.6 μL drop at 12 mg/mL of the complex of ARL15 (32-197) and CNNM2 CBS domain (429-584) in HPLC buffer mixed with 0.6 μL of reservoir solution containing 0.2 M sodium chloride, 0.1 M Tris pH 8.5, 25% (w/v) PEG3350. Crystals grew in 1-2 days at 20 ºC. For data collection, crystals were cryo-protected by soaking in the reservoir solution supplemented with 30% (v/v) ethylene glycol.

### Structure solution and refinement

Diffraction data from single crystals of ARL15 (32-197)-CNNM2 CBS complex were collected at the Advanced Photon Source (APS) (Supp. Table X). Data processing and scaling were performed with DIALS (Otwinowski and Minor, 1997). The initial phases for the complex structure were determined by molecular replacement with Phaser (McCoy et al., 2007), using the coordinates of the CNNM2 CBS domain (PDB entry 4IY0) (Gimenez-Mascarell *et al*., 2017) and an AlphaFold model of ARL15 (Senior et al., 2020). The initial phases were improved by Autobuilder in PHENIX package (Adams et al., 2010). The starting protein model was then completed and adjusted with the program Coot (Emsley and Cowtan, 2004) and improved by multiple cycles of refinement, using the program phenix.refine (Adams *et al*., 2010) and model refitting. At the latest stage of refinement for both structures, we also applied the translation-libration-screw (TLS) option (Winn et al., 2003). The final models have 99.5% residues in the allowed regions of Ramachandran plot. The coordinates have been deposited with the Protein Data Bank (PDB) under the accession number 8F6D. Refinement statistics are given in Table 1.

**Table 1.** Statistics of data collection and refinement

| PDB code | 8F6D |
| --- | --- |
| **Data collection** |  |
| X-ray source | APS 24ID-E |
| Wavelength (Å) | 0.9792 |
| Space group | P1 |
| Cell dimensions |  |
| *a*, *b*, *c* (Å) | 66.02, 73.32, 79.33 |
| $\alpha$, $\beta$, $\gamma$ (°) | 94.4, 95.5, 115.3 |
| Resolution (Å) | 50-3.20 (3.26-3.20)<sup>1</sup> |
| *R*<sub>merge</sub> | 0.078 (0.782) |
| *I* / $\sigma I$ | 10.4 (0.6) |
| Completeness (%) | 94.2 (93.1) |
| Redundancy | 3.7 (3.8) |
| CC<sub>1/2</sub> | 0.998 (0.624) |
| **Refinement** |  |
| Resolution (Å) | 29.5 - 3.20 |
| No. reflections | 20526 |
| *R*<sub>work</sub> / *R*<sub>free</sub> | 0.264/0.292 |
| No. atoms |  |
| Protein | 7791 |
| *B*-factors |  |
| Protein | 115.3 |
| R.m.s deviations |  |
| Bond lengths (Å) | 0.004 |
| Bond angles (°) | 0.79 |
| Ramachandran statistics (%) |  |
| Most favored regions | 98.0 |
| Additional allowed regions | 1.6 |
| Disallowed regions | 0.4 |

<sup>1</sup>Highest resolution shell is shown in parentheses.

### GST-pulldown assays

For GST-pulldown with purified recombinant proteins, a slurry of 25 μl of Glutathione Sepharose beads (Cytiva Sweden AB) was washed twice with 1 ml of pulldown buffer (50 mM HEPES pH 7.5, 200 mM NaCl, 1 mM MgCl_2_, 1 mM TCEP-HCl, 0.02% Igepal). In between the washes, beads were sedimented by centrifuging at 13,000 rpm for 1 min at 4 °C and supernatant was discarded. 200 μl of 1 mg/ml GST-fused protein was added to the beads and incubated on ice for 15 mins. The beads were washed twice again with 1 ml of buffer as mentioned previously. 50 μl of the binding partner (1 mg/ml) was added to the beads and incubated on ice for 30 min. On washing the beads thrice with 200 μl of buffer, the proteins were eluted with 25 μl of 20 mM reduced glutathione solution. 20 μl of the eluate was transferred and mixed with 5 μl of 5 × SDS loading dye. 15 μl of the sample was loaded onto the gels. Pulldown results were verified by SDS-PAGE. For GST-pulldown of ARL15 and mutants expressed in HEK-293T cells, GST and GST-fused CBS-pair domain of murine CNNM2 were expressed in transformed BL21 (DE3) cells (Stratagene, CA). Bacterial cells were lysed by sonication in ice-cold PBS containing 1% Triton X-100 and protease inhibitor phenylmethylsulfonyl fluoride (Sigma–Aldrich). The bacterial cell lysates were then incubated with glutathione agarose (Sigma–Aldrich) overnight at 4 ºC with rotation. The agarose beads were washed with PBS with 1% Triton X-100. Concentrations of the GST-fused proteins were measured by Coomassie stain on the SDS–PAGE gel using serial diluted BSA proteins as controls. To express ARL15 proteins, 10 μg of the wild-type and mutant ARL15 plasmids were transiently transfected into HEK-293T cells plated in a 10-cm dish. After 24 hours, the cells were lysed in 1 ml of mild lysis buffer containing protease inhibitor mixture (Roche Life Sciences) and phosphatase inhibitor mixture (EMD Millipore), and spun down at 14,000 × *g* for 10 min at 4 °C. For GST-pulldown assays, the cell lysate supernatants were incubated with glutathione agarose bound with 20 μg of GST or GST-fused CBS-pair domain proteins overnight at 4 ºC with rotation.

The bound proteins were washed with 1 ml of PBS with 0.1% Triton X-100 three times by rotation and eluted into 50 μl of SDS sample buffer. Lysates input (20 μl) and pulldown samples were separated on SDS–PAGE gels and analyzed by immunoblotting. The rabbit monoclonal FLAG antibody (#14793) was used to detect expressed FLAG-tagged ARL15 proteins. SDS-PAGE and Coomassie staining was employed to visualize and demonstrate equivalent amount of GST and GST-fused CBS-pair domain in the assay. SDS-PAGE and western blot was used to demonstrate equivalent amount of FLAG-tagged ARL15 wild-type and mutants as inputs in the pulldown assay. SDS-PAGE and western blot also used to analyze the binding of FLAG-tagged ARL15 wild-type and mutants to GST compared to GST-fused CBS-pair domain.

### Cell Lines

Cultured human cells were maintained in a Dulbecco’s Modified Eagle Medium (DMEM), high glucose media with 10% FBS in a humidified 37 °C, 5% CO_2_ incubator. The 293T cell line, here referred as HEK-293T cells, was purchased from ATCC (CRL-3216, Manassas, Virginia, USA). 293-TRPM7 cells, expressing FLAG-tagged mouse TRPM7 under tetracycline control, was generously provided by Dr. Andrew Scharenberg (University of Washington).

### Co-immunoprecipitation

A 10 cm dish of HEK-293T cells were transfected with 8 μg of FLAG-tagged human ARL15 wild-type (WT) or the FLAG-tagged ARL15 R95A mutant (R95A) using the Turbofect Transfection Reagent. All the following biochemical procedures were conducted at 4 ºC. 24 hours post-transfection cells were lysed with 800μl of mild lysis buffer (50mM TRIS pH 7.4, 150 mM NaCl, 1% Igepal 630) containing protease inhibitors. Proteins were solubilized by incubating the lysis mixture for 30 min. The samples were then cleared by centrifugation at 15,600 × g for 10 min. Supernatants (lysates) were subjected to immunoprecipitation as follows. FLAG-tagged ARL15 proteins were immunoprecipitated using 50 μL of Pierce Anti-DYKDDDDK Magnetic Agarose (ThermoFisher Scientific) for 2 hours at 4 ºC. The beads were washed two times with PBS and once with purified water. The bound proteins were eluted with 50μL of 2X Laemmli sample buffer, and the proteins resolved by SDS-PAGE and Western Blotting using standard procedures. The anti-FLAG M2 antibody (Sigma-Aldrich) was used to detect FLAG-tagged ARL15 proteins. The anti-CNNM3 antibody (NBP2-32134, Novus Biologicals) was used to detect endogenous human CNNM3.

### Electrophysiological recordings

The voltage-clamp technique was used to evaluate the whole-cell currents of TRPM7 expressed in 293-TRPM7 cells as previously described (Li et al., 2007). Briefly, whole-cell current recordings of TRPM7-expressing cells were elicited by voltage stimuli lasting 250 ms delivered every 1 sec using voltage ramps from –100 to +100 mV. Data were digitized at 2 or 5 kHz and digitally filtered offline at 1 kHz. The internal pipette solution for macroscopic current recordings contained 145 mM Cs methanesulfonate, 8 mM NaCl, 10 mM EGTA, and 10 mM HEPES, pH adjusted to 7.2 with CsOH. The extracellular solution for whole-cell recordings contained 140 mM NaCl, 5 mM KCl, 2 mM CaCl_2_, 10 mM HEPES, and 10 mM glucose, pH adjusted to 7.4 (NaOH).

### Zn^2+^ influx assay

The Zn^2+^ influx assay used to characterize TRPM7 channel function has been previously described in detail (Bai *et al*., 2021). Briefly, cells were plated into 24-well dishes coated with polylysine to aid comparison between individual samples by fluorescence microscopy. Before labeling, cells were washed with Hanks’ balanced salt solution (HBSS) containing 0.137 M NaCl, 5.4 mM KCl, 0.25 mM Na_2_HPO_4_, 6 mM glucose, 0.44 mM KH_2_PO_4_, 1.3 mM CaCl_2_, 1.0 mM MgSO_4_, and 4.2 mM NaHCO_3_. Cells were then labeled with the Zn^2+^ indicator FluoZin-3 (2.5 μM) in HBSS following manufacturer instructions (Thermo Fisher Scientific). The cells were then washed once with HBSS and then placed back into HBSS before images were quickly acquired on an inverted Olympus IX70 fluorescence microscope with a 10X phase contrast objective (Olympus Neoplan 10/0.25 Ph, Olympus, Tokyo, Japan). Cells were visually inspected for uneven dye loading prior to imaging. To stimulate Zn^2+^ influx, 30 μM ZnCl_2_ in HBSS was introduced to cells. For static measurements, images of cells from the different samples were taken at a specific time point 5 to 10 minutes after the addition of 30 μM ZnCl_2_.

## Acknowledgements

We thank Dr. Alexei Gorelik for crystallographic data collection and assistance with data processing. X-ray data were acquired at the Advanced Photon Source, a U.S. Department of Energy, Office of Science user facility operated by Argonne National Laboratory under Contract No. DE-AC02-06CH11357.

## Supplemental materials

**Supplemental Figure S1.**
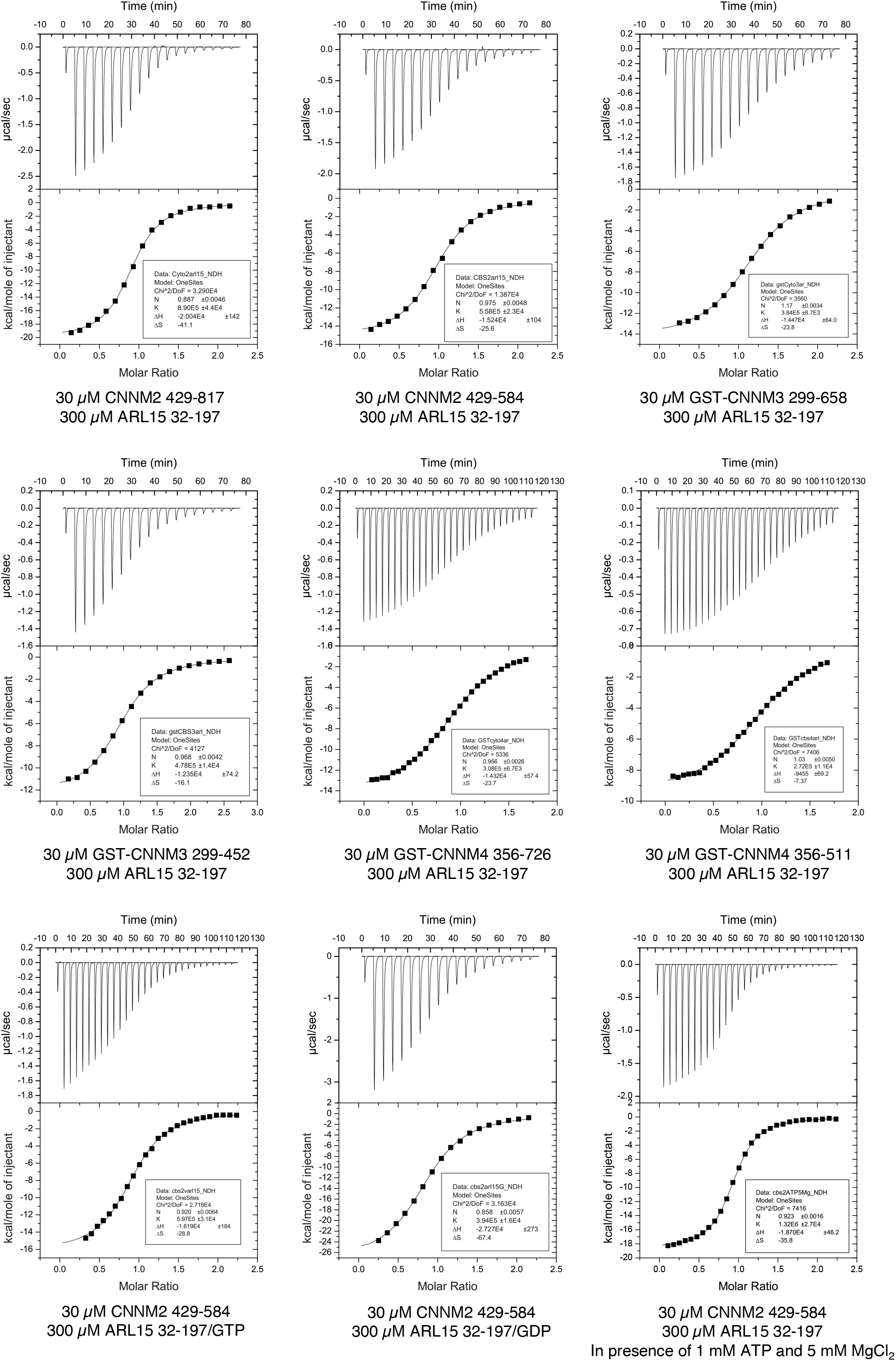
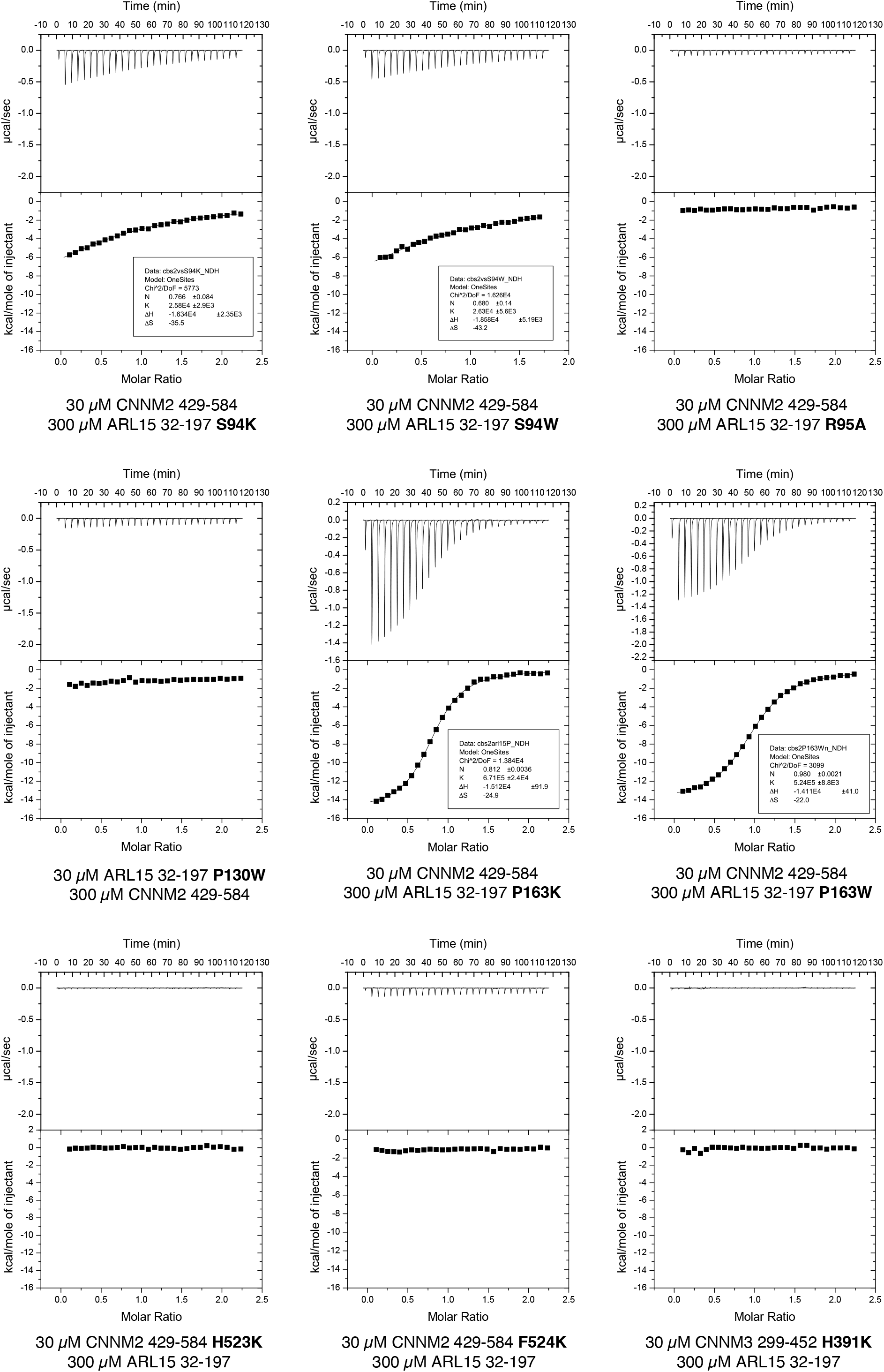
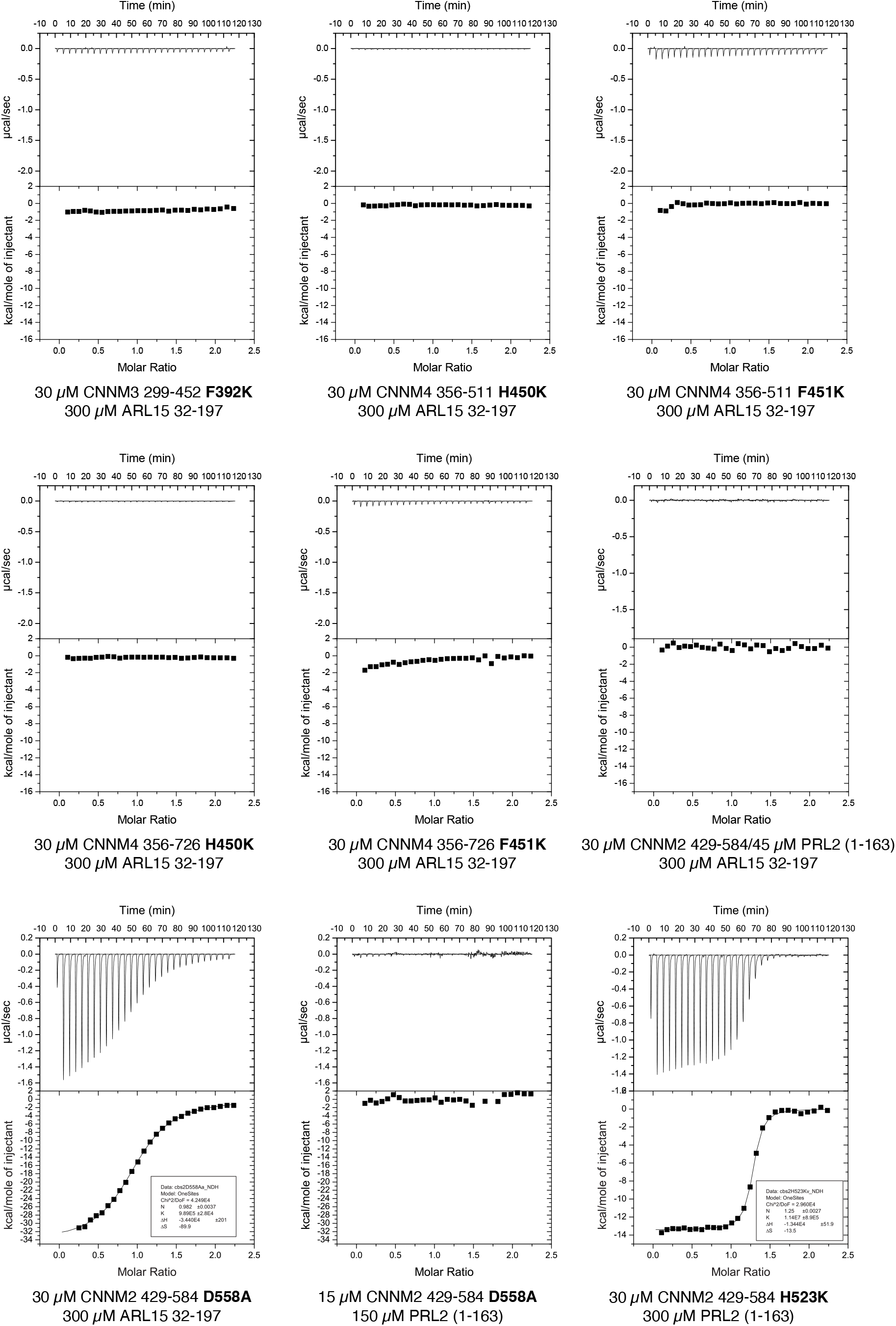
ITC thermograms. Protein concentrations in the cell and the syringe are indicated.

**Supplemental Figure S2.**
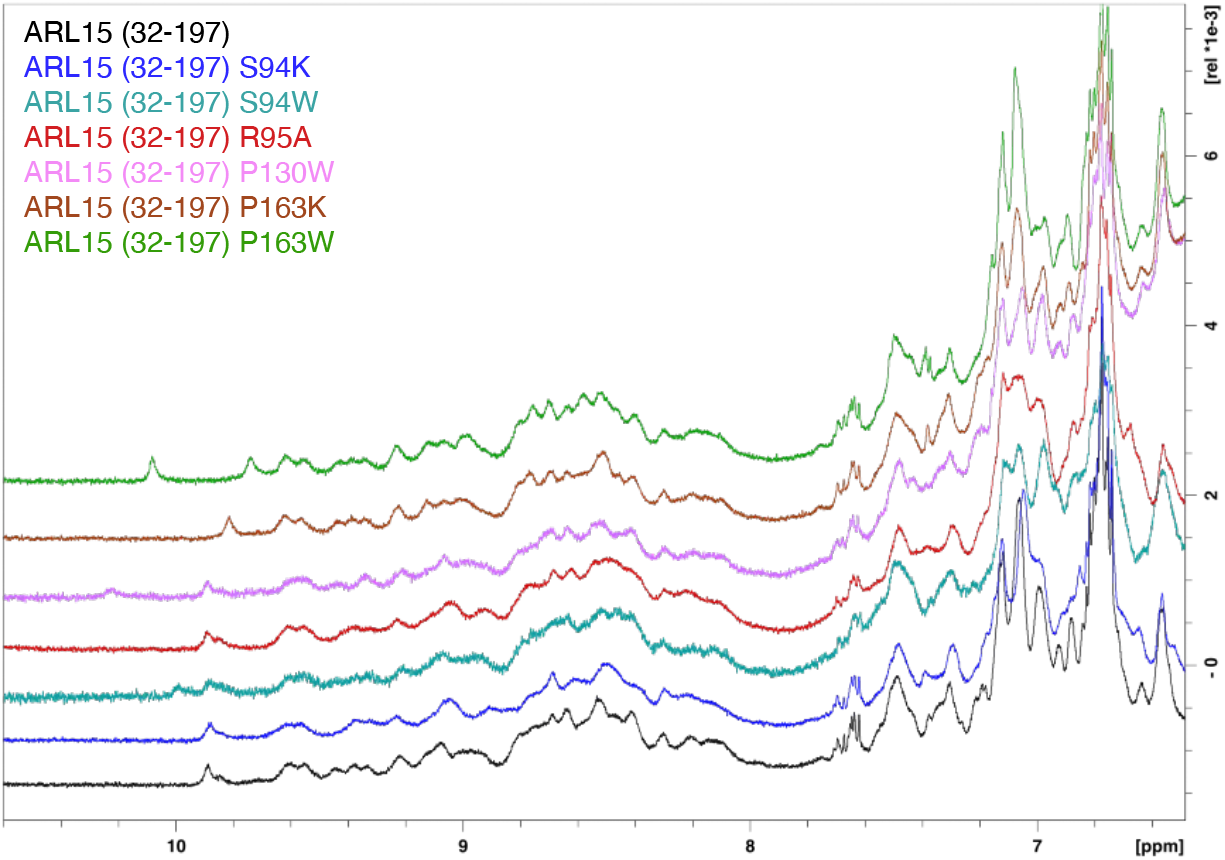
Downfield ^1^H NMR spectra of ARL15 GTPase domain (residue 32-197) and its mutants (S94K, S94W, R95A, P130W, P163K, P163W).

**Supplemental Figure S3.**
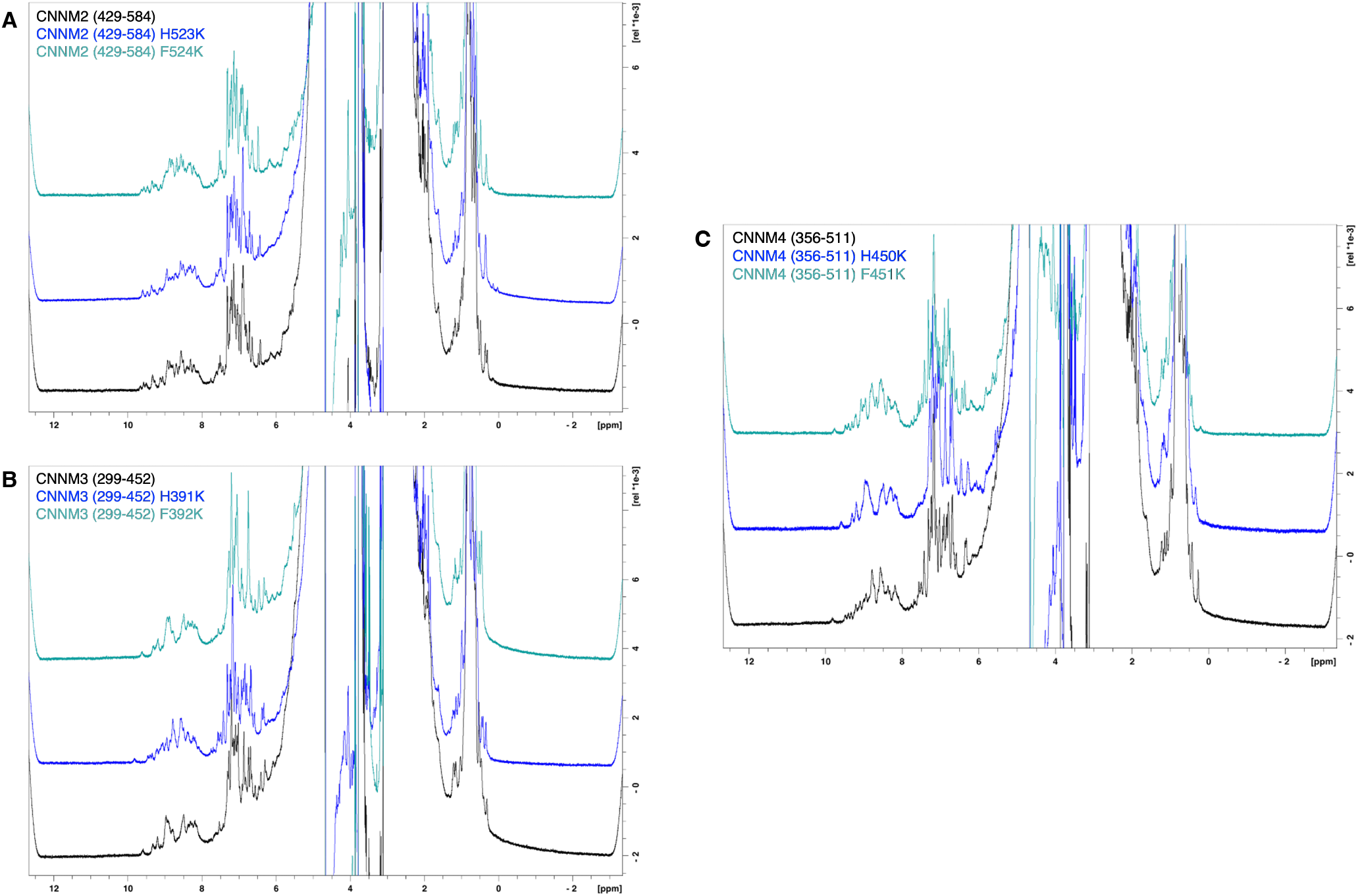
^1^H NMR spectra of CNNM CBS-pair domains and their mutants. **A**, CNNM2 (429-584), CNNM2 (429-584) H523K, CNNM2 (429-584) F524K. **B**, CNNM3 (299-452), CNNM3 (299-452) H391K, CNNM3 (299-452) F392K. **C**, CNNM4 (356-511), CNNM4 (356-511) H450K, CNNM4 (356-511) F451K.

**Supplemental Figure S4.**
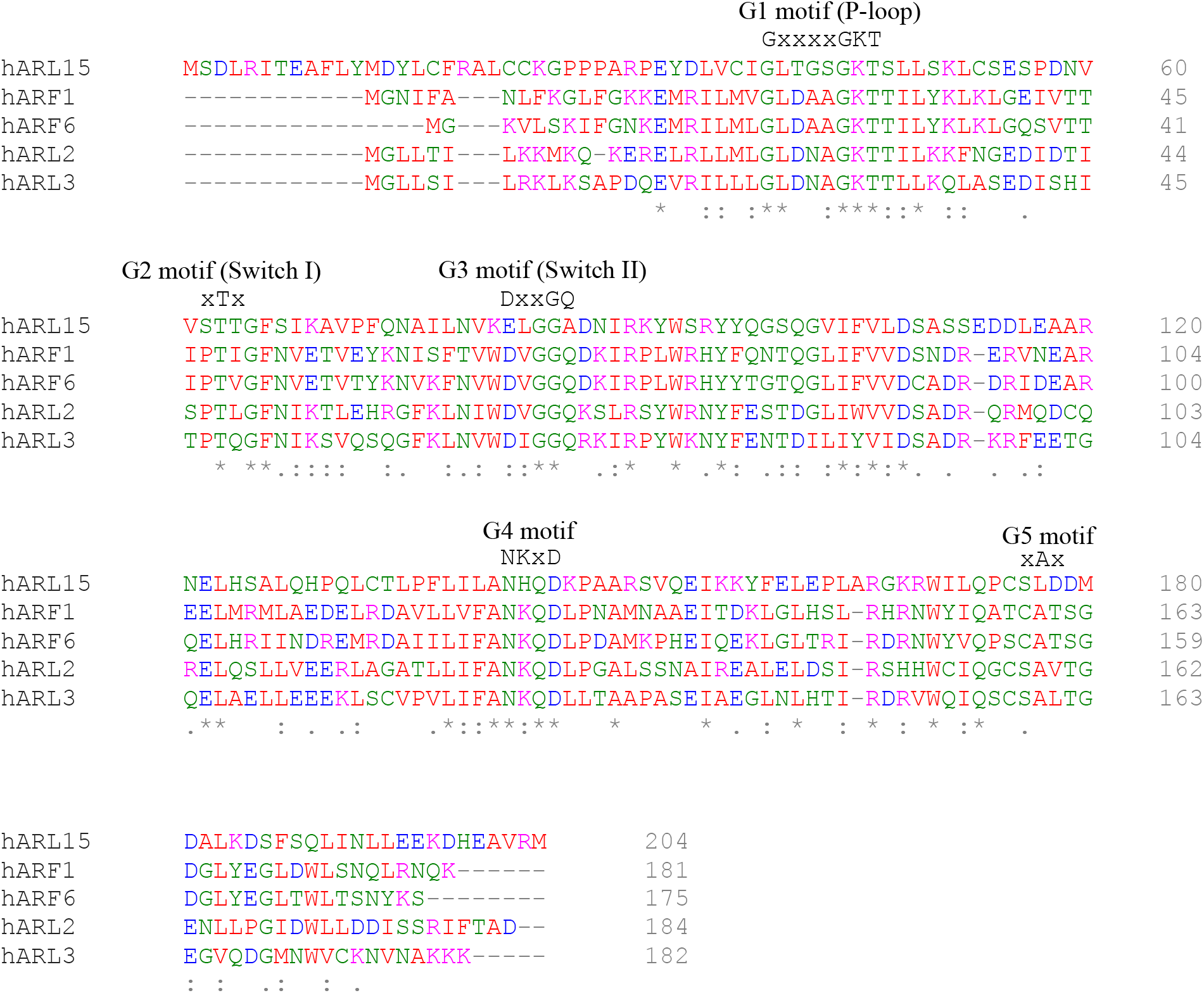
Sequence alignment of human ARL15 and other ARL/ARF GTPases. The glutamine residue that is unique to ARL15 is in the G3 motif.

